# Structural Diversity and Phylogenetic Distribution of Valyl tRNA-like Structures in Viruses

**DOI:** 10.1101/2020.06.19.162263

**Authors:** Madeline E. Sherlock, Erik W. Hartwick, Andrea MacFadden, Jeffrey S. Kieft

## Abstract

Viruses commonly use specifically folded RNA elements that interact with both host and viral proteins to perform functions important for diverse viral processes. Examples are found at the 3′ termini of certain positive-sense ssRNA virus genomes where they partially mimic tRNAs, including being aminoacylated by host cell enzymes. Valine-accepting tRNA-like structures (TLS^Val^) are an example that share some clear homology to canonical tRNAs but have several important structural differences. Although many examples of TLS^Val^ have been identified, we lacked a full understanding of their structural diversity and phylogenetic distribution. To address this, we undertook an in-depth bioinformatic and biochemical investigation of these RNAs, guided by recent high-resolution structures of a TLS^Val^. We cataloged many new examples in plant-infecting viruses but also in unrelated insect-specific viruses. Using biochemical and structural approaches, we verified the secondary structure of representative TLS^Val^ substrates and tested their ability to be valylated, finding structural heterogeneity within this class. In a few cases, large stem-loop structures are inserted within distinct variable regions located in an area of the TLS distal to known host cell factor binding sites. In addition, we identified one virus whose TLS has switched its anticodon away from valine; the implications of this remain unclear. These results refine our understanding of the structural and functional mechanistic details of tRNA mimicry and how this may be used in viral infection.

## INTRODUCTION

RNAs adopt complex structures to accomplish a wide range of functions and often the same class of RNA appears in a wide variety of species but with substantial variation. For example, within a given class of RNA the primary sequence may vary, but conservation of base pair patterns and specific nucleotides preserve key tertiary interactions or form critical functional elements. For some RNAs the overall global shape is not critical as long as a conserved functional core exists within the larger fold (e.g. to bind ligands or perform catalysis) (Doherty and Doudna 2001; Weinberg et al. 2015; McCown et al. 2017), while other RNAs rely almost entirely on their overall three-dimensional shape (e.g. to block a processive nuclease or mimic the structure of another RNA) (Pfingsten et al. 2006; Dreher 2010; Chapman et al. 2014; Akiyama et al. 2016; Pisareva et al. 2018). Finally, it is possible for different classes of RNAs with distinct secondary and tertiary structures to perform identical functions (Corbino et al. 2005; Perreault et al. 2011; Steckelberg et al. 2018a). Understanding the fundamental rules of the RNA structure-function relationship requires detailed explorations of diverse RNAs across and within functional classes.

An interesting class of functional RNA elements found in viruses are the transfer RNA (tRNA)-like structures (TLSs) found at the 3′ terminus of certain positive-sense single-stranded plant-infecting RNA viruses (Pinck et al. 1970; Yot et al. 1970; Rietveld et al. 1983; Mans et al. 1991; Dreher 2010). These RNAs were identified by their ability to induce *in cis* aminoacylation of the 3′ end of the viral RNAs by host cell aminoacyl tRNA synthetases (AARSs) (Pinck et al. 1970; Yot et al. 1970; Hall et al. 1972; Öberg and Philipson 1972). The implied structural mimicry makes them intriguing examples of RNA-based molecular mimicry (Rietveld et al. 1983; Hammond et al. 2009). The three known classes of TLSs have distinct secondary structures compared to one another and to canonical tRNAs, and each class is charged with a different amino acid: valine, histidine, or tyrosine (Mans et al. 1991; Hammond et al. 2009; Dreher 2010). TLSs play multiple roles that confer an advantage during infection (Hall 1979; Haenni et al. 1982; Mans et al. 1991; Dreher 2009; Dreher 2010), including *in cis* enhancement of translation of the viral RNA (Matsuda and Dreher 2004). While some portions of these viral TLSs resemble a tRNA, they are typically larger, and their positions at the 3′ end of the viral RNA mandate an alternate connectivity involving a 3′ pseudoknot structure and the D-loop at the 5′ end that then leads to the remainder of the viral genome, whereas a tRNA’s 5′ and 3′ ends are paired in its acceptor stem (Rietveld et al. 1982; Rietveld et al. 1983; Pleij et al. 1985; Felden et al. 1994).

Valine-accepting TLSs (TLS^Val^) are on average the smallest of the three classes and share the most homology to tRNAs (Dreher and Goodwin 1998). This homology allows TLS^val^ to interact with many of the same host factors as tRNAs including the CCA-adding enzyme, valyl-tRNA synthetase (ValRS) and eukaryotic elongation factor 1A (eEF1A) (Pinck et al. 1970; Yot et al. 1970; Giege et al. 1978; Joshi et al. 1982; Matsuda and Dreher 2004). In addition, the TLS contains the promotor for viral negative-strand synthesis and therefore also interacts with the viral RNA-dependent RNA polymerase (RdRp) (Deiman et al. 1998; Singh and Dreher 1998). The structure of the prototype TLS^Val^ from turnip yellow mosaic virus (TYMV) has been studied for decades using a variety of biochemical and biophysical techniques (Rietveld et al. 1982; Matsuda and Dreher 2004; Hammond et al. 2009; Hammond et al. 2010), but only recently has high-resolution structural information become available. Specifically, two structures of the TYMV TLS^Val^, both on its own (Colussi et al. 2014) or attached to its 5′ upstream pseudoknot domain (UPD) (Hartwick et al. 2018), were solved by X-ray crystallography. These structures revealed fundamental similarities and differences between the TYMV TLS^Val^ and tRNAs, identified key tertiary contacts needed to stabilize the fold, and showed how the TLS and UPD domains are oriented and how they communicate.

The structures of the TYMV TLS raise new questions about the TLS^Val^ class as a whole, and also enable new explorations. Specifically, while several dozen TLS^Val^ are known in plant-infecting viruses (Dreher and Goodwin 1998; Goodwin and Dreher 1998; Dreher 2010), we do not have a full catalog of this type of tRNA mimicry. However, secondary and tertiary structural information from high-resolution structures (Colussi et al. 2014; Hartwick et al. 2018) can now be combined with new search tools to find additional examples. Because some features may be idiosyncratic, analyses of new examples help to understand the diversity within a class and therefore how different elements form tertiary interactions or contacts with viral or host factors. Comparative sequence analysis using homology searches is a powerful tool to expand knowledge of RNA classes such as riboswitches, ribozymes, exoribonuclease-resistant RNAs, and others of unknown function (Barrick et al. 2004; Weinberg et al. 2007; Weinberg et al. 2010; Roth et al. 2014; Weinberg et al. 2015; Weinberg et al. 2017; Steckelberg et al. 2018b). Therefore, we employed structure guided bioinformatic strategies to identify new examples of RNAs conforming to the TLS^Val^ pattern. These used primary sequence and secondary structure conservation guided by information from three-dimensional crystal structures to iteratively search large databases for new sequences. Using chemical probing and functional assays, we verified putative novel examples and explored the extent of their conservation and diversity. Our findings reveal previously unrecognized structural and potentially functional variations in the TLS^Val^ RNA class, including the verification of TLS^Val^ within insect-infecting viruses.

## RESULTS AND DISCUSSION

### Homology searches reveal additional putative examples of valyl tRNA-like structures

We conducted homology-based searches starting with a seed alignment containing 28 previously identified examples of TLS^Val^ from the Rfam database (http://rfam.xfam.org/family/RF00233) (Kalvari et al. 2018). We made adjustments to this initial alignment based on the crystal structure of the TYMV TLS (Colussi et al. 2014) and previously proposed models for other members of this class (Goodwin and Dreher 1998). Homology searches using the program Infernal (Nawrocki and Eddy 2013) identified 108 unique sequences of putative TLS^Val^ RNAs in 46 distinct viruses (Fig. 1, see also Supplemental Files 1 and 2). Some of these viruses contain multiple RNA segments, which often contain similar but not identical putative TLS^Val^ sequences at their 3′ ends.

**FIGURE 1.**
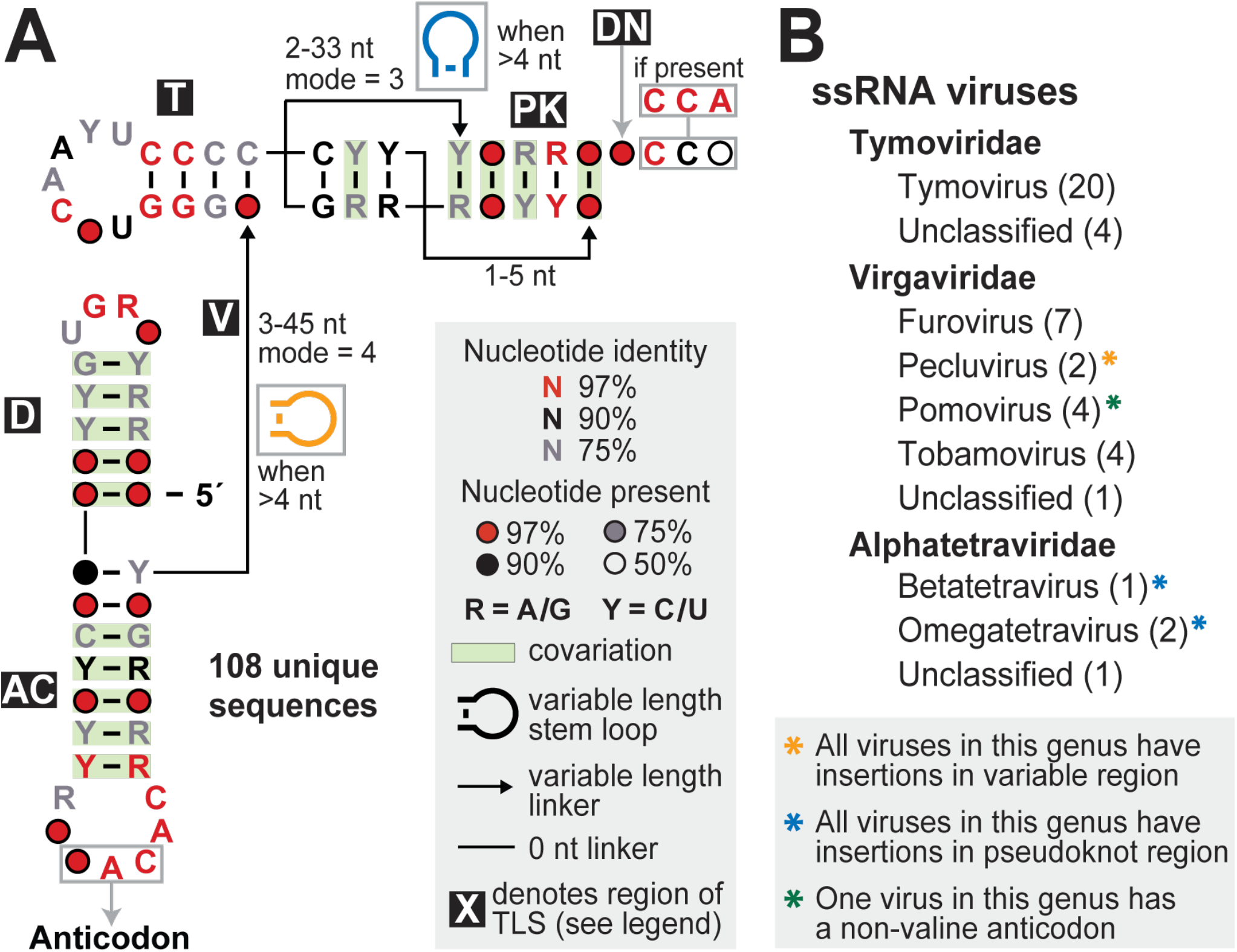
Structural conservation and phylogenetic distribution of all valyl tRNA-like structures in viruses. (*A*) Consensus sequence and secondary structure model of the 108 unique examples of valine-accepting tRNA-like structures. Each portion of the structure is labeled, mostly according to its homolog in a canonical tRNA (D: D-arm; AC: anticodon arm; V: variable region; T: T-arm; PK: pseudoknot region; DN: discriminator nucleotide). Stem-loop structure insertions are sometimes present in the linkers within the variable and pseudoknot regions. (*B*) Genera that contain valine-accepting tRNA-like structures, further organized by family. The number of individual viruses that contain these structures within each genus is listed in parentheses and all examples are derived from positive-sense single-stranded RNA viruses.

The compiled unique sequences were used in R-scape (Rivas et al. 2017; Rivas et al. 2020) to calculate a consensus sequence and secondary structure model, which has several key features (Fig. 1A). First, all of the base-paired elements are supported by covarying mutations except the part homologous to the tRNA’s T-arm. The TLS^Val^ T-loop is well conserved, to a similar extent as a canonical tRNA, but the stem is always capped by two non-covarying GC pairs. While typically four base pairs in length, some examples may contain only those two GC base pairs or use non-canonical pairing at the base of the T-arm stem. Second, except for three examples discussed below, the anticodon always encodes valine (generally ‘NAC’) and the wobble position can be any of the four nucleotides. Third, within a set of tRNAs specific for a given amino acid, the ‘discriminator nucleotide’ located just upstream of the 3′ terminal CCA is conserved (Crothers et al. 1972; Hou 1997). In contrast, in TLS^Val^ the discriminator nucleotide is not significantly conserved (Fig. 1A) although often it is an A, matching tRNA^Val^. Finally, because these viral genomes do not encode a terminal A, the viral genome is 3′-adenylated by the host cell’s CCA-adding enzyme (Giege et al. 1978). This, and the technical limitation that many sequencing reads are truncated, causes the nucleotides at the 3′ terminus to be under-represented in the TLS^Val^ alignments. Therefore, while the terminal ‘CCA’ does not appear to be significantly conserved, when present they are 100% conserved (see Supplemental File 1 for all aligned sequences).

Most of the putative TLS^Val^ RNAs are found in RNAs of the *Tymoviridae* family with several other examples in the *Virgaviridae* family (Fig. 1B). It has previously been noted that TLS^Val^ is more abundant in *Tymoviridae* and TLS^His^ is more abundant in *Virgaviridae* but each is occasionally found in the other family (Dreher 2010). We also identified four putative TLS^Val^s in insect-infecting tetraviruses, which are positive sense ssRNA viruses that are often segmented. It was previously proposed that the 3′ terminus of both Nudaurelia capensis beta virus (NCBV) and Helicoverpa armigera Stunt Tetravirus (HaSV), two of the four tetravirus examples, form a tRNA-like structure (Gordon et al. 1995; Gordon et al. 1999), but these RNAs were not tested for aminoacylation and the proposed secondary structure model differed from known TLS^Val^. Specifically, these models (Gordon et al. 1995; Gordon et al. 1999) proposed 5′ to 3′-end base pairing as in tRNAs instead of the 3′ pseudoknot arrangement that has since been demonstrated to form by other members of TLS^Val^. Overall, most new putative examples of TLS^Val^ are predicted to closely resemble the prototypic TYMV, but a number contain variations including insertions in several linker regions as well as differences in the anticodon identity. These observations, and the presence of putative TLS^Val^ in insect-infecting viruses, are explored and discussed here.

### Most TLS^Val^s conform to a shared secondary structure

Secondary structure predictions based on conservation across species provide compelling evidence for conserved stems and pseudoknots within a class of RNA, but it is critical to experimentally evaluate the structure of individual sequence representatives to ensure proper alignment. This is especially true in areas that are too highly conserved for covarying base pairs to be observed or in sites where substantial insertions are predicted. Thus, we performed in vitro chemical probing experiments for several different TLS^Val^ RNAs. The relative reactivity of each nucleotide as determined by selective 2′ hydroxyl acylation analyzed by primer extension (SHAPE) probing experiments resolved by capillary electrophoresis (Yoon et al. 2011; Kim et al. 2013; Cordero et al. 2014; Kladwang et al. 2014; Lee et al. 2015) was determined for each TLS^Val^ RNA and mapped onto the corresponding secondary structure prediction (Fig. 2A-B; Fig S1). Using the prototype TYMV TLS as a positive control, we observed higher reactivity in known loops and variable regions and relatively low reactivity in base-paired elements, consistent with previous results and the crystal structure (Hartwick et al. 2018) (Fig. S1A). We then probed another representative TLS^Val^ from Japanese soil-born wheat mosaic virus (JSWMV), which conforms well to the consensus model. Probing data from this putative TLS^Val^ matched TYMV, indicating the same overall fold (Fig. S1B). These results strongly suggest that putative TLS^Val^s that conform well to the consensus model share a common secondary structure and likely similar tertiary structures.

**FIGURE 2.**
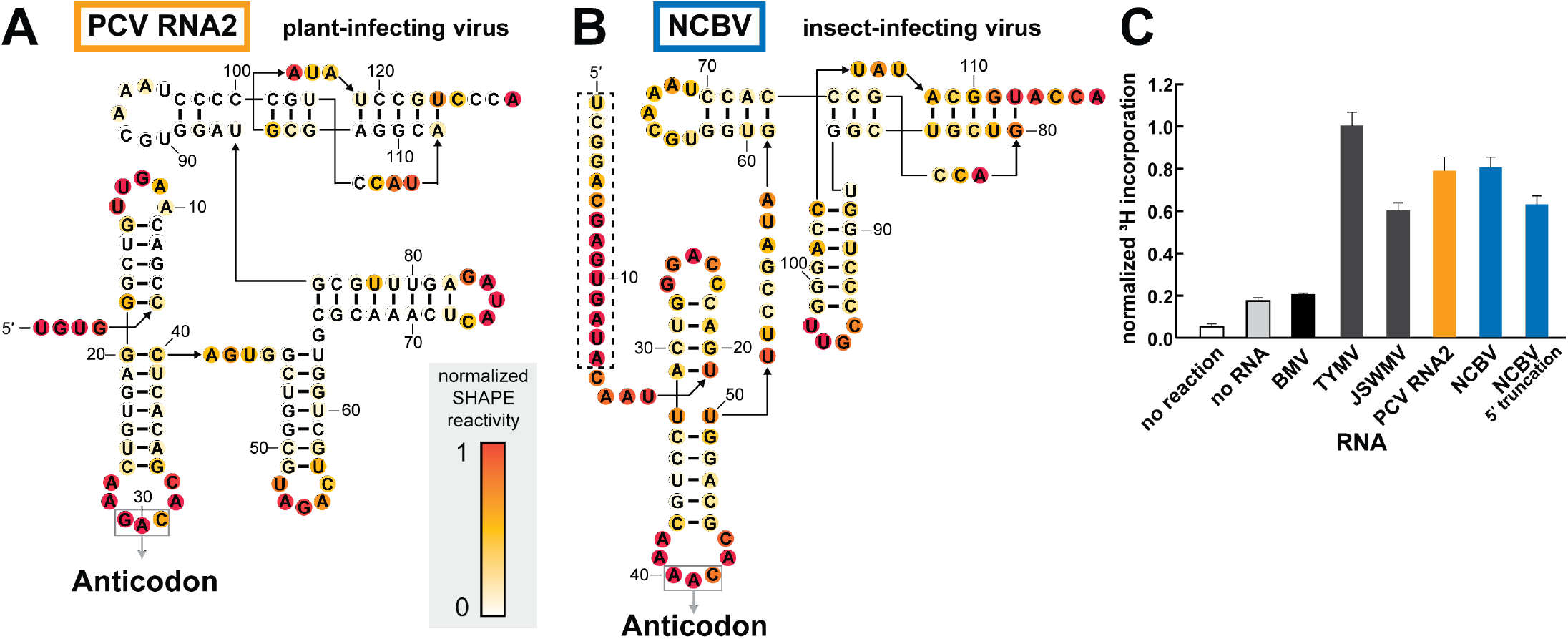
Divergent TLS^Val^ RNAs containing stem-loop insertions are competent substrates for valylation. (*A, B*) Chemical probing of TLS representatives from Peanut clump virus (A) and Nudaurelia capensis beta virus (B) using the SHAPE reagent NMIA. Reactivity was background subtracted and normalized according to the reactivity of loop regions in hairpin structures (not shown) flanking the TLS structure on both the 5′ and 3′ ends. (*C*) Activity of valyl tRNA synthetase (ValRS) on TLS RNAs as measured by the covalent addition of radiolabeled (^3^H) valine at their 3′ termini. The ^3^H incorporation, as measured by a scintillation counter, was normalized to the TYMV TLS^Val^ construct, which had been previously tested and optimized under these reaction conditions. The truncated NCBV construct begins at nucleotide C15 (see panel B for sequence). The BMV TLS belongs to a separate class of tyrosine-accepting TLSs. Orange and blue bars correspond to the color schemes for PCV and NCBV, respectively, in panels A and B as well as in Figure 4.

### Substantial insertions are tolerated in discrete locations within TLS^Val^

The main source of structural heterogeneity in the TLS^Val^ is predicted insertions in the parts of the TLS that correspond to the “variable” region between the anti-codon arm and T-arm, or within a loop of the 3′ acceptor stem pseudoknot (Fig. 1A). The former is found in examples from peanut clump virus (PCV) RNAs, and the secondary structures predicted by our bioinformatic searches agree with a previously proposed model that places two large stem-loop structures in the variable region (Goodwin and Dreher 1998). The latter occurs in three of the four examples derived from insect-infecting Tetraviruses, but in those our predicted secondary structures were different than in previously published models (Gordon et al. 1995; Gordon et al. 1999).

To determine the correct secondary structure of these divergent putative TLS^Val^s, we applied chemical probing to RNAs from PCV and NCBV as representative of plant-infecting and insect-infecting viruses. The reactivity patterns were consistent with the bioinformatic predictions of an overall secondary structure pattern that matches the “typical” TLS^Val^, but with stem-loops inserted into the aforementioned regions (Fig. 2A-B). Because the NCBV RNA was previously proposed to have a different secondary structure, we mapped the reactivity data on both the predicted structure from this study (3′ pseudoknot model) and the previously proposed 5′ to 3′ pairing model (Gordon et al. 1999). The pattern is consistent with our new model proposed by the bioinformatic searches but not the previous model (Fig. S1C).

### Divergent TLS^Val^ RNAs are aminoacylated in vitro

We then asked if the divergent TLS^Val^s could serve as substrates for aminoacylation. While the TLS^Val^ RNAs from plant-infecting TYMV, JSWMV, and PCV had all been shown to undergo valylation *in vitro* (Dreher and Goodwin 1998; Goodwin and Dreher 1998), putative examples from insect-infecting viruses had never been tested. We tested all of these RNAs for *in vitro* valylation using recombinantly purified ValRS and ^3^H-labeled valine under conditions previously used for the TYMV TLS (Hartwick et al. 2018). Negative control reactions containing no RNA or the TLS^Tyr^ from Brome mosaic virus (BMV) show low levels of valylation. However, all TLS^Val^ RNAs, including NCBV, show valine incorporation levels significantly above background (Fig. 2C). Thus, aminoacylatable TLS^Val^ RNAs exist in insect-infecting viruses.

We used the ability of the NCBV TLS^Val^ to be valylated to further interrogate its bioinformatically predicted structural model (3′ pseudoknot) versus the previously reported model (5′ to 3′ paired). Because the 5′ to 3′ paired model includes ~15 nucleotides not involved in predicted structures in the 3′ pseudoknot model (Fig. S1C) (Gordon et al. 1999), we tested a truncated version that cannot form the 5′ to 3′ pairing (Fig. 2B). This 5′ truncated RNA is still valylated, which likely would not occur if the 5′ to 3′ paired model were valid. Note that ^3^H-valine incorporation is lower for the truncated construct as compared to the extended construct, which could be due to the absence of some optional substructures analogous to the TYMV TLS when the upstream pseudoknot domain is included (Dreher and Goodwin 1998; Hammond et al. 2010).

Taken together, our results indicate that large insertions in some TLS^Val^s are structurally and functionally tolerated. Whether these confer additional function is unknown. Our results also show that TLSs are not exclusively a plant virus phenomenon but are also found in animal-infecting viruses. While our bioinformatic searches only revealed four examples of this, additional divergent TLS^Val^ examples likely remain to be found in other viruses, and/or completely unrelated TLS classes may be discovered in other species.

### TLSs with non-valine anticodons fail to be aminoacylated

The bioinformatic analysis identified three putative TLS^Val^s with a non-valine anticodon in the three RNA segments of Colombian potato soil-borne virus (CPSbV), all of which have a UAA (leucine) in the anticodon loop (Gil et al. 2016). The TLS sequences from these three segments of CPSbV are similar (Fig. S2), but a few covarying mutations in base-paired regions and one other mutation in a variable region make the strict conservation of the UAA anticodon peculiar. A sequencing error seems unlikely since the UAA leucine anticodon was present in all of the RNAs in multiple isolates of this recently-identified virus (Gil et al. 2016). This anticodon triplet is only one nucleotide away from a valine anticodon (UAC is valine-compatible) yet none of the three examples contain a valine codon, raising the question of why all three RNA segments have converted to the same non-valine anticodon.

To verify the alignment that assigned the UAA triplet as the anticodon, we performed chemical probing of the putative TLS RNA from CPSbV. The pattern was consistent with the secondary structure model for TLS^Val^ and supported that the UAA was properly placed, creating a leucine anticodon (Fig. 3A). We then tested the sequence from CPSbV RNA3 for *in vitro* valylation. Consistent with the non-valine anticodon, the CPSbV RNA3 was not aminoacylated (Fig. 3B), nor were RNA1 or RNA2 (see Supplemental File 2). However, mutation of the anticodon from UAA (leucine) to CAC (valine) in CPSbV RNA3 conferred the ability of this RNA to be valylated (Fig. 3B), hence other than their anticodons these RNAs have retained all other TLS^Val^ features.

**FIGURE 3.**
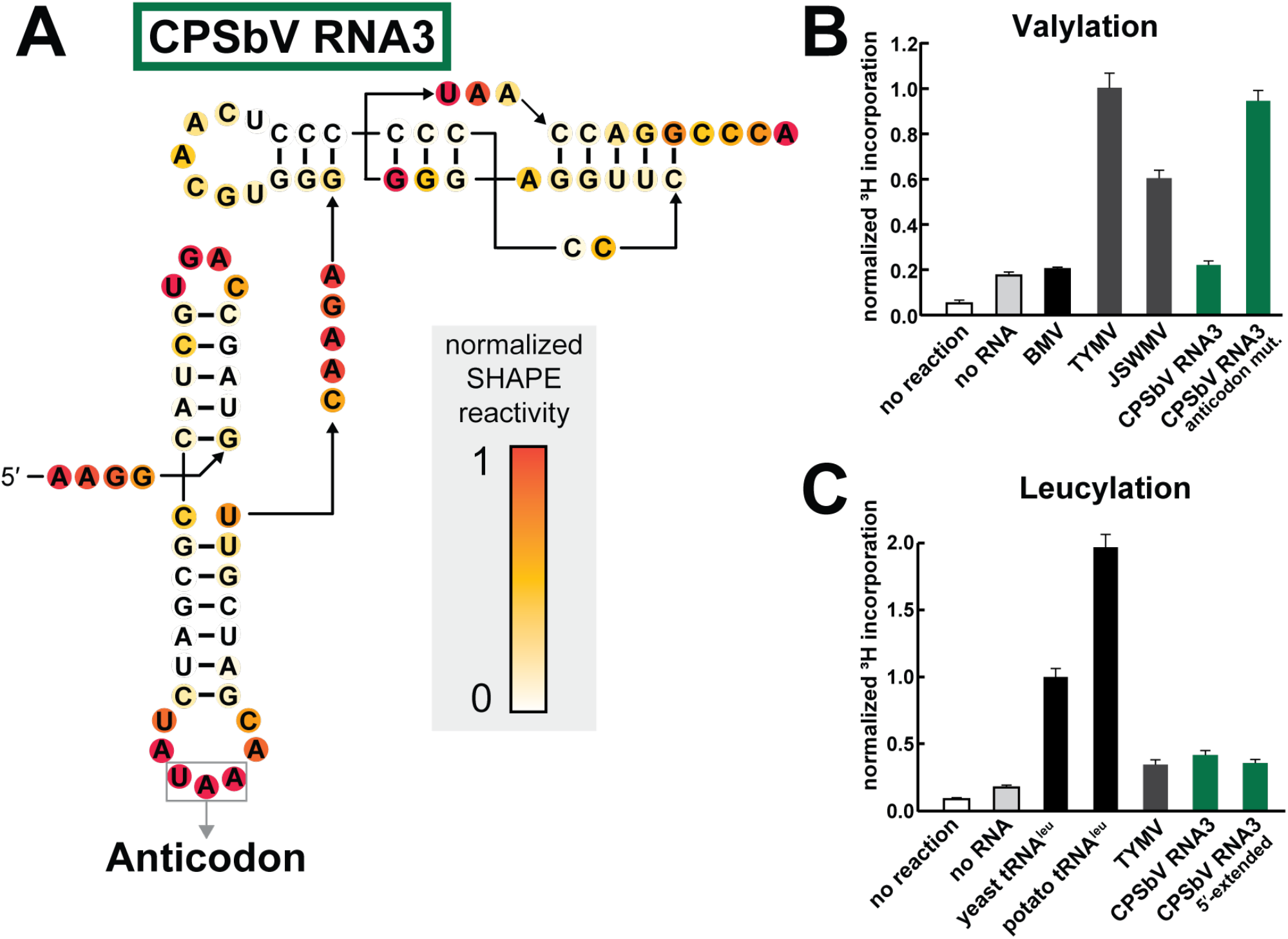
The leucine anticodon of the CPSbV TLS prevents in vitro valylation activity. (*A*) Chemical probing of TLS representatives from Colombian potato soil-borne virus using the SHAPE reagent NMIA. Additional annotations and details are described in the legend to Fig. 2. (*B*) Activity of valyl tRNA synthetase (ValRS) on CPSbV TLS RNAs as measured by the covalent addition of radiolabeled (^3^H) leucine at their 3′ termini. The WT sequence for CPSbV TLSs corresponds to a leucine anticodon, as depicted in panel A, while the anticodon mutation construct contains two mutations (UAA→CAC) that alter the anticodon identity to valine. For additional details see Fig. 2B. (*C*) Activity of leucyl tRNA synthetase (LeuRS) on TLS RNAs or tRNAs as measured by the covalent addition of radiolabeled (^3^H) leucine at their 3′ termini. Leucylation was measured by scintillation counter to determine ^3^H-Leucine incorporation and normalized to the yeast tRNA^Leu^ construct. The CPSbV 5′-extended construct includes an additional 65 nucleotides upstream of the sequence depicted in panel A (see Table S1 for sequence details). Green bars correspond to the color scheme in panels A and B as well as in Figure S2.

It seemed possible that CPSbV RNAs might indicate a new class of TLSs with an overall secondary structure that conforms to the TLS^Val^ class, but which are charged with leucine by the cognate synthetase. We tested this using recombinant leucyl-tRNA synthetase (LeuRS) from *Saccharomyces cerevisiae* and ^3^H-leucine. To verify the activity of the *S. cerevisiae* LeuRS enzyme we tested its ability to aminoacylate positive control in vitro transcribed tRNA^Leu^ from *S. cerevisiae* and tRNA^Leu^ from *Solanum tuberosum,* the virus’ potato plant host (Fig. 3C). In vitro transcribed TYMV TLS^Val^ served as the negative control RNA and was not efficiently aminoacylated. Testing a CPSbV TLS, we did not find any conditions under which it was aminoacylated with leucine above the negative control (Fig. 3C). We then tested another CPSbV RNA3 construct with an additional 65 nucleotides upstream of the beginning of the TLS structure. This includes the portion of the 3′ UTR conserved across all three genomic RNAs, as we speculated that additional structures in this region could be necessary for or increase the efficiency of leucylation. However, the extended CPSbV RNA also is not leucylated above the negative control. Variation of buffer conditions, RNA concentration, enzyme concentration, or length and temperature of incubation (see Methods for details) did not yield leucine incorporation signal above the negative control (see Supplemental File 2).

The inability of the *S. cerevisiae* LeuRS to aminoacylate the CPSbV TLS could be due to a high degree of specificity for the AARS of the host species *S. tuberosum*. While we cannot rule out this possibility, it seems unlikely given the robust leucylation signal measured from a *S. tuberosum* tRNA^Leu^ by the yeast enzyme. Furthermore, previous structural and functional characterization of LeuRS show that this synthetase does not appear to query the anticodon (Fukunaga and Yokoyama 2005b). Modeling the TYMV TLS into a complex with the LeuRS enzyme suggests that the anticodon would not be recognized (Figure S3). Rather, LeuRS recognition of tRNA^Leu^ relies on other elements (Fukunaga and Yokoyama 2005a; Fukunaga and Yokoyama 2005b) that are absent from the putative CPSbV TLS. Specifically, a stem-loop in the variable region of tRNA^Leu^ is recognized by the LeuRS but the CPSbV TLS lacks this. Additionally, the discriminator nucleotide for tRNA^Leu^ is an A that is important for recognition by LeuRS (Fukunaga and Yokoyama 2005a) but all three CPSbV TLS RNAs contain a C at this position. Given these requirements, it is perhaps not surprising that the putative CPSbV TLSs would not be recognized by the LeuRS.

Why would the anticodon in the TLS of all three CPSbV RNAs specifically encode a leucine if the rest of the TLS structure matches a valine tRNA, especially if this prevents any aminoacylation? While we cannot exclude the possibility that the CPSbV TLSs are leucylated in the context of a cell and therefore represent a separate class from the rest of the TLS^Val^ RNAs, our preliminary assessment is that this is unlikely. It is possible these are indeed valine-accepting TLSs, but our in vitro aminoacylation assays fail to fully mimic the host cell environment, and we cannot rule out that the *S. tuberosum* ValRS is able to valylate these RNAs in spite of their leucine anticodon. Perhaps the CPSbV TLSs are a poorer substrate than the other TLS^Val^ RNAs and it is beneficial for CPSbV to be aminoacylated inefficiently or not at all. There are a number of viruses related to those with TLS^Val^ that lack substantial portions of the canonical TLS^Val^ structure needed for aminoacylation and therefore do not have an amino acid added to the 3′ end of the genomic RNAs (Dreher and Goodwin 1998). However, in the case of CPSbV a small substitution mutation converts them to true TLS^Val^, and it is odd that evolution would alter all three copies in such a targeted way if selection is just to eliminate aminoacylation. Indeed, this pattern suggests that these TLSs could readily convert to authentic TLS^Val^. Maybe it is advantageous for the virus to have a specific species of uncharged tRNA-like element on its 3′ end, perhaps activating stress-response pathways that in some way favor viral proliferation. Our data are not able to address these possibilities, but they hint at an intricate interplay between viral RNA and host machinery in CPSbV infection that will require infection-based experiments to explore.

### The consensus model suggests protein binding strategies are conserved

Previous superpositions of the TYMV TLS^Val^ structure onto the structure of an authentic tRNA^Val^ ValRS complex and a tRNA bound to EF-Tu showed how it could bind those factors, as the overall geometry and key surfaces of a tRNA are maintained in the TYMV TLS (Colussi et al. 2014). Superimposing the TYMV TLS+UPD structure onto these portions likewise shows the UPD does not interfere with these interactions (Fig. 4) or with binding of the CCA-adding enzyme (Fig. S4). Examination of the consensus model suggests that all TLS^Val^s can make the same binding contacts. Specifically, the 3’ end acceptor stem pseudoknot is present and twelve base pairs are maintained from the T-loop to just before the 3ʹ CCA motif, as observed in tRNA and the crystallized TYMV crystal structures (Colussi et al. 2014; Hartwick et al. 2018). Thus, the length of the T-loop-acceptor stem helical stack is maintained across the class. In the tRNA^Val^-ValRS structure, the anticodon loop is highly distorted (Fukai et al. 2000) in order to be recognized by the synthetase. While the anticodon loop of the TYMV TLS does not adopt this conformation in the RNA-only crystal structure, it was previously hypothesized that the loop is flexible enough to adopt the same conformation as that of tRNA^Val^ when bound by ValRS (Colussi et al. 2014). The anticodon loop is highly conserved among TLS^Val^ RNAs and it is likely that all members of the class are recognized in this manner by ValRS. Put together, the consensus model is consistent with evolution constraining the structures to maintain features essential for interactions with ValRS, eEF1A, and the CCA-adding enzyme.

**FIGURE 4.**
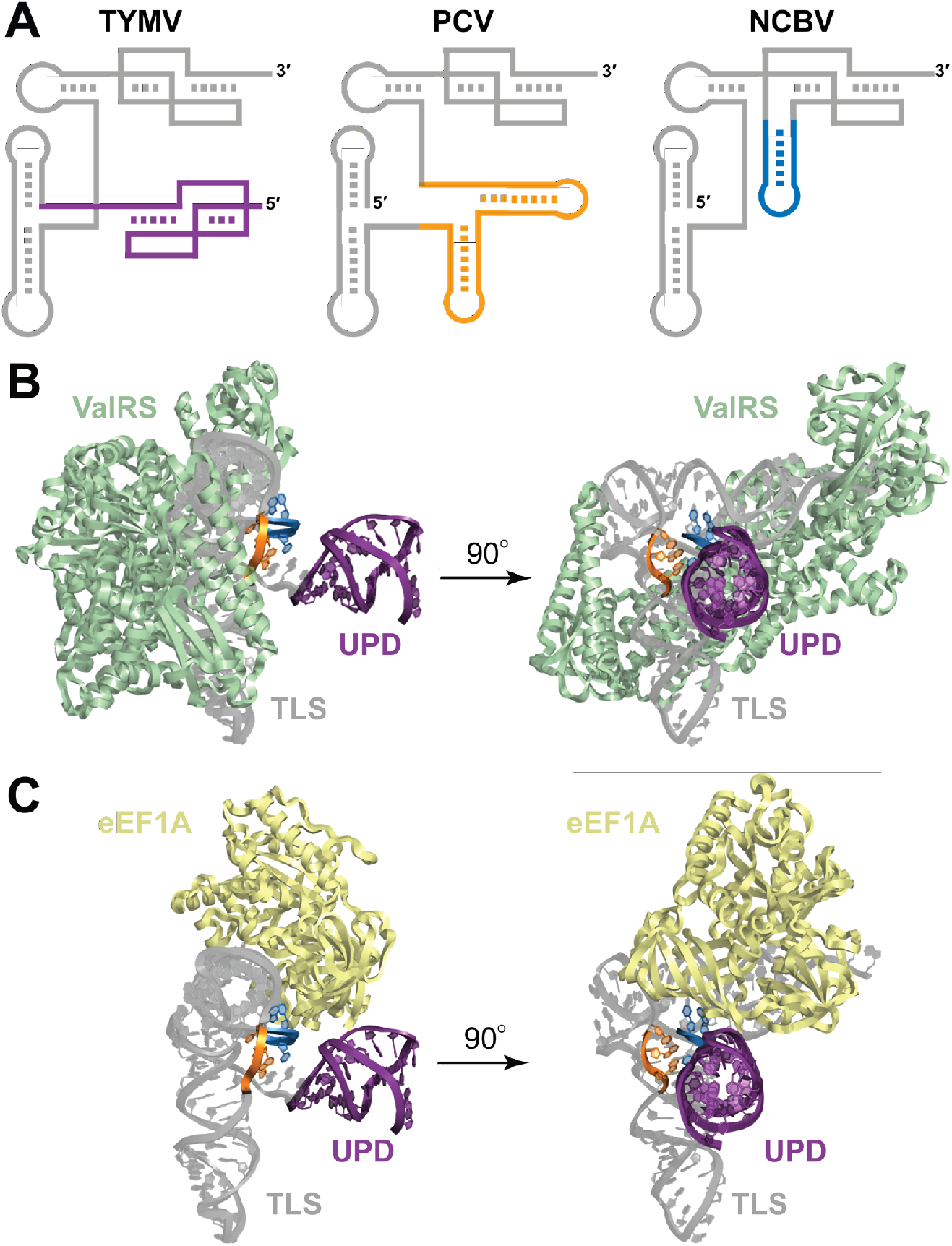
Stem-loop insertions found in divergent TLS^Val^s do not interfere with host factor interactions. (*A*) Cartoon models of the TLS representatives from TYMV, PCV, and NCBV. The upstream pseudoknot domain (UPD) of the TYMV TLS is drawn in purple (note that the PCV and NCBV TLSs are not preceded by a UPD in the context of the viral genomic RNA). The stems inserted within the variable region of the PCV TLS are orange and the insertion within the 3’ pseudoknot structure of NCBV is blue. (*B*) Modeling of the TYMV TLS (gray) with the valyl-tRNA synthetase (ValRS; green) from *Thermus thermophilus*. The nucleotides corresponding to the site of insertion for PCV (variable region) and NCBV (3ʹ pseudoknot) are colored orange and blue, respectively, and the UPD is colored purple to match panel (A). (*C*) Modeling of the TYMV TLS (gray) with eEF1A (yellow) from *Oryctoagus cuniculus*. Colors are as for panel (B).

### Insertions are positioned to not interfere with protein binding

The presence of substantial stem-loop insertions in several TLS^Val^s (Fig. 4A) are tolerated in terms of aminoacylation (Fig. 2) and thus do not seem to interfere with ValRS binding. To correlate this with structure, we mapped the location of these insertions onto the model of the TYMV TLS+UPD on ValRS and on eIF1A. Both insertion points lie on the face of the TYMV TLS where the UPD and genome connect and thus not where they will interfere with synthetase, eIF1A or CCA-adding enzyme binding (Fig. 4, Fig. S4). Interestingly, the insertion points are very close to where the UPD is attached to the TYMV TLS^Val^, suggesting that the stem-loop insertions would clash with a UPD if one were present. However, we find no strong evidence that the PCV or NCBV TLS^Val^s have UPDs 5′ of their predicted TLS regions and thus the inserted stem-loops can apparently occupy the same three-dimensional space as the UPD in TYMV. It is tempting to speculate that convergent evolution has resulted in several ways to place an RNA structural domain in this location to perform a similar function, but this remains untested.

### The identity of the nucleotide at the discriminator position reflects the viral family

The discriminator nucleotide for tRNA^Val^ is nearly always an A, while in the TLS^Val^ it is not well conserved but is usually an A or sometimes a C. The ValRS-tRNA^Val^ complex structure (Fukai et al. 2000) suggests no obvious interactions with the discriminator nucleobase and thus it could be a purine or pyrimidine. In the TLS^Val^ this nucleotide may play a more significant role in interactions with a viral protein factor than a host factor. Indeed, the stem of the 3′ pseudoknot, the discriminator nucleotide, and the terminal CCA are important sites of recognition for the TYMV RNA-dependent RNA polymerase (RdRp) (Deiman et al. 1998). Nearly all TLS^Val^ RNAs with an A at the discriminator nucleotide position are of the *Tymoviridae* family, while a C is almost exclusively found in the *Virgaviridae* family. Consistent with this observation, the TYMV RdRp strongly prefers a purine nucleotide in this position (Deiman et al. 1998), while the RdRp from tobacco mosaic virus (TMV), a prototypical *Virgaviridae* member, is most efficient with a C at this position (Osman et al. 2000). Hence, variations in the TLS^Val^ evolve to match the specific needs of the virus, while maintaining the ability to interact with important host factors.

### Concluding remarks

Consensus models based on the full set of RNAs of a given class help give a full understanding of the key characteristics necessary for function, contextualize the characteristics of each member, reveal the presence of interesting structural or functional variations, and suggest how evolution can fine-tune structure. Here, we built on the list of known TLS^Val^s and also verified the presence and structure of interesting variants, including those with insertions, altered anticodons and in insect-infecting viruses. These discoveries open new questions regarding changes in function or advantage to the virus conferred by these differences as well as guiding studies aimed at understanding the mechanisms of events such as translation enhancement and RdRp recruitment. Finally, observing the variations seen here underscores the possibility that other classes of tRNA-like structures outside of the known TLS^Val^, TLS^His^, or TLS^Tyr^ remain to be identified, with architectures sufficiently different as to not appear in homology searches.

## METHODS

### Bioinformatic Searches

A preliminary alignment of all known TYMV-like TLSs (Rfam ID: RF00233) was obtained from the Rfam database (Kalvari et al. 2018). Additional examples of this class of RNAs were identified using Infernal version 1.1 (Nawrocki and Eddy 2013) to query a compilation of all viral sequence reads in the National Center for Biotechnology Information (NCBI) nucleotide database (downloaded 01/22/2019). Iterative searches were performed and adjustments were made to the alignment based on information from the high resolution crystal structure of the TYMV TLS (Colussi et al. 2014; Hartwick et al. 2018) as well as previously reported secondary structure models for the PCV TLS RNAs (Goodwin and Dreher 1998). Duplicate sequences were removed, resulting in 108 unique sequences derived from 46 unique viruses. The consensus sequence and secondary structure model as well as the statistical analysis of covariation were calculated using the RNA Significant Covariation Above Phylogenetic Expectation (R-scape) program (Rivas et al. 2017; Rivas et al. 2020) visualized using R2R (Weinberg and Breaker 2011) and labeled in Adobe Illustrator.

### RNA Preparation

DNA templates were produced either by PCR amplification from a double-stranded DNA gene block fragment or by combining two overlapping single-stranded DNA oligonucleotides and using a SuperScript II Reverse Transcriptase (Invitrogen) reaction, according to manufacturer protocol, to fill in the remaining template. All DNA oligonucleotides used in this study were purchased from Integrated DNA technologies (IDT) and sequences can be found in Table S1. Reverse primers used to amplify the constructs for in vitro aminoacylation assays contained two 5ʹ-terminal 2ʹOMe-modified bases to achieve a higher yield with the correct 3’ end of the construct, since the 3ʹ terminal adenosine nucleotide is the site of modification. All RNA oligonucleotides used in this study were produced by *in vitro* transcription using templates produced by PCR or RT. Each 200 μL transcription reaction contained: 30 mM Tris pH 8.0, 60 mM MgCl_2_, 8 mM each NTP, 10 mM DTT, 0.1% spermidine, 0.1% Triton X-100, and T7 RNA polymerase. Transcription reactions were incubated at 37 °C overnight then purified by denaturing 10% PAGE. Gels were visualized by UV illumination and the gel piece containing the band of correct length was excised. Gel pieces were sliced into small pieces and soaked in ~300 μL of diethylpyrocarbonate (DEPC)-treated milli-Q filtered water (Millipore) water at 4 °C overnight to elute the RNA. The supernatant containing RNA was subjected to ethanol precipitation then resuspended in DEPC-treated water and diluted to the appropriate concentration.

### Protein expression and purification

The ValRS enzyme from *T. thermophilus* used in the current study was recombinantly expressed and purified previously and details can be found in the Methods section of reference: (Hartwick et al. 2018). The LeuRS enzyme from *S. cerevisiae* was purified for this study. LeuRS Gene Blocks (IDT) were cloned into a pET15b vector using Gibson ligation. The protein was expressed in BL21 (DE3) cells in LB containing ampicillin at 37 °C until OD_600_ = 0.2 – 0.3, the temperature was reduced to 18 °C and continued to grow until the OD_600_ = 0.6, then was induced with 0.25 mM IPTG overnight. The cells were harvested and spun at 5488 x g for 12 minutes at 6 °C. The pellets were resuspended in lysis buffer containing 50 mM Tris-HCl pH 8, 500 mM NaCl, 2 mM β-mercaptoethanol, 5 mM MgCl_2_, 10% glycerol, and 1 mM EDTA-free protease inhibitor tablet (Roche) then sonicated for 2 min total processing time (20 sec on, 40 sec off). The lysate was centrifuged at 30,000 x g for 30 min at 4 °C before being loaded into a gravity flow column (Bio-Rad) and purified by Ni-NTA resin (UBP Bio) followed by size exclusion using FPLC and a Sepax 300 SEC column. The protein was resuspended to 2 μM and stored in buffer containing 50 mM Tris-HCl pH 8, 5 mM MgCl_2_, 2 mM β-mercaptoethanol, and 5% glycerol.

### Chemical probing of RNAs *in vitro*

Structure probing experiments using the chemical modifier *N*-methyl isatoic anhydride (NMIA) were performed according to the previously published protocol from reference: (Cordero et al. 2014). Briefly, RNAs were refolded by heating to 90 °C for 3 minutes, cooled to room temperature, then modified by incubating for 20 minutes at room temperature with either NMIA or DMSO (final concentrations: 60 nM RNA, 3 mg/mL NMIA or DMSO, 50 mM HEPES-KOH pH 8.0, 10 mM MgCl_2_, 3 nM FAM-labeled DNA primer for reverse transcription, see Table S1 for sequence). The probing reaction was quenched by adding NaCl (final concentration 500 mM) and Na-MES buffer (final concentration 50 mM, pH 6.0) and oligo-dT magnetic beads (Invitrogen Poly(A)Purist MAG Kit). Chemically modified RNAs were purified using the magnetic stand and washed with 70% ethanol. Reactions were resuspended in water, then reverse transcription was performed using SuperScript III enzyme (Invitrogen). RNA ladders were produced by four separate reverse transcription reactions using ddNTPs. Reverse transcription reactions were incubated at 50 °C for 45 minutes, then the RNA was degraded by adding NaOH (final concentration 200 mM), heating to 90 °C for 5 minutes, then quenching with an acidic solution (final concentration: 250 mM sodium acetate pH 5.2, 250 mM HCl, 500 mM NaCl). The remaining DNA products were purified using the magnetic stand then washed with 70% ethanol. A solution containing HiDi formamide solution (ThermoFisher) and spiked with GeneScan 350 ROX Dye Size Standard (ThermoFisher) was added to elute DNA products from the magnetic beads. Labeled RT DNA products were analyzed by capillary electrophoresis using an Applied Biosystems 3500 XL system. Fragment size analysis, alignment, background subtraction, and normalization (based on reactivity in flanking stem-loop regions) were performed using the HiTrace RiboKit (https://ribokit.github.io/HiTRACE/) (Yoon et al. 2011; Kim et al. 2013; Kladwang et al. 2014; Lee et al. 2015) with MatLab (MathWorks), and figures were produced using RiboPaint (https://ribokit.github.io/RiboPaint/tutorial/) with Matlab subsequently labeled in Adobe Illustrator.

### In vitro aminoacylation assays using ^3^H-labeled amino acids

In vitro-transcribed RNAs were resuspended in DEPC-treated water to 1 μM and refolded by incubating at 90 °C for 3 minutes then cooling to room temperature. Aminoacylation reactions were set up by mixing 1 μL of RNA or water, 1 μL of buffer (10X: 300 mM HEPES-KOH, pH 7.5, 20 mM ATP, 300 mM KCl, 50 mM MgCl_2_, 50 mM dithiothreitol), 1 μL ^3^H-labeled L-valine (60 Ci/mmol) or ^3^H-labeled L-leucine (100 Ci/mmol), 1 μL of ValRS or LeuRS enzyme (10X: 2 μM) and 6 uL of water (final volume = 10 μL). In additional experiments that attempted to optimize leucylation signal for the CPSbV TLS, the following conditions were altered: increased RNA concentration (5 or 10 μM RNA instead of 1 μM 10X stock), increased Mg^2+^ and ATP concentration (10X: 40 mM ATP, 100 mM MgCl_2_), or completely altered buffer conditions according to previous studies (Goodwin and Dreher 1998): IV buffer (10X: 25 mM Tris-HCl pH 8.0, 20 mM MgCl_2_, 10 mM ATP, 1 mM spermine) or TM buffer (10X: 300 mM HEPES-KOH pH 7.5, 1M potassium acetate, 25 mM magnesium acetate, 15 mM ATP). Aminoacylation reactions, each performed in triplicate, were incubated at 30 °C for 2-3 hours then immediately loaded onto a vacuum filter blotting apparatus. Filtering was achieved by using one layer each, ordered from bottom to top, of thick filter paper (BioRad gel dryer filter paper), HyBond positively charged membrane (GE Healthcare) and 0.45 μm Tuffryn membrane filter paper (PALL Life Sciences) and each layer was pre-washed and equilibrated with a wash buffer (20 mM Bis-Tris pH 6.5, 10 mM NaCl, 1 mM MgCl_2_). After blotting, each reaction blot was washed five times with 200 uL of wash buffer that included trace xylene cyanol for visualization. After filtering and washing, the filters were dried, then the Hybond membrane was cut out, placed in a scintillation vial, and ^3^H incorporation was measured, taking two readings per sample, by a scintillation counter (Perkin-Elmer Tri-Carb 2910 TR). Data were analyzed and plotted using Microsoft Excel.

### Modeling

A composite model of the TYMV 3ʹ UTR using both crystal structures for the complete TLS and UPD domains (PDB IDs: 4p5j (Colussi et al. 2014) and 6mj0 (Hartwick et al. 2018), respectively) was built using coot (Emsley et al. 2010) as previously performed (Hartwick et al. 2018). Modeling and coloring of the TYMV 3ʹ UTR was performed using PyMOL (Schrodinger 2015) and Protein Data Bank (Berman et al. 2000; Burley et al. 2019) (PDB: rscb.org) deposited structures for ValRS (Fukai et al. 2000) (PDB ID: 1gax), eEF1A (Shao et al. 2016) (PDB ID: 5lzs), LeuRS (Fukunaga and Yokoyama 2005b) (PDB ID: 1wz2), and archaeal CCA-adding enzyme (Kuhn et al. 2015) (PDB ID: 4×4r). In each case, the TLS model was aligned to best match the tRNA present in each structure using PyMOL.

## ACKNOWLEDGEMENTS

We thank members of the Kieft Lab for many discussions and David Constantino and Dr. Quentin Vicens for critical reading of this manuscript. We thank Dr. Steve Bonilla Rosales for assistance with aminoacylation assays. This work was support by NIH grant R35GM118070 to JSK. MES is a Jane Coffin Childs Postdoctoral Fellow. EWH was a University of Colorado School of Medicine RNA Bioscience Initiative Scholar.

